# Chemical quantification of elemental sulfur identifies S_8_ as a redox-active sulfur metabolite

**DOI:** 10.1101/2025.09.26.678775

**Authors:** Nada E. Eldakra, Ethan J. York, Rebecca L. Cordell, Christopher H. Switzer

## Abstract

Elemental sulfur occupies a central position in biological sulfur chemistry as a thermodynamically stable endpoint of reactive sulfur species (RSS) interconversion, yet its role in biology has remained poorly defined due to the lack of robust analytical methods for its direct measurement. Here, we describe a selective, accessible chemical workflow for the detection and quantification of zero-valent elemental sulfur (S^0^) and elemental selenium (Se^0^) in complex biological systems, enabling systematic interrogation of these species as biological redox metabolites. Elemental sulfur is isolated from biological samples as the cyclo-octasulfur (S_8_) allotrope via phase-selective organic solvent extraction and structurally confirmed by Raman spectroscopy. Extracted S_8_ is subsequently derivatized with triphenylphosphine (TPP) to form triphenylphosphine sulfide (TPP=S), which is quantified using complementary ^31^P–nuclear magnetic resonance (NMR) spectroscopy and GC–MS, providing sensitivity across nanomolar to millimolar concentration ranges. The method exhibits strong chemical selectivity against competing sulfur species and is compatible with a wide range of biological matrices, including conditioned media, cultured cells, tissues, and solid biological materials. Extension of the workflow to selenium chemistry enables selective detection and quantification of elemental selenium (Se^0^) following chemical or cellular reduction of selenite. Collectively, these results establish a practical and transferable analytical platform that renders zero-valent chalcogen species experimentally accessible in biological contexts. By enabling direct measurement of elemental sulfur across diverse biological systems, this method reveals S^0^ as a previously inaccessible, redox-active sulfur metabolite and provides a foundation for defining its formation, distribution, and function in cellular and tissue redox biology.

**Highlights:**

∘ S^0^ and Se^0^ are predicted redox endpoints, yet selective detection is limited.
∘ A broadly accessible chemical method is introduced to quantify S^0^ and Se^0^.
∘ Quantification spans nM–mM ranges in diverse matrices, *e*.*g*., cells and media.
∘ Reveals S^0^ as a redox-active metabolite present in multiple biological matrices.

## Introduction

Reactive sulfur species (RSS) are abundant in biological systems and arise from central sulfur metabolism, redox reactions, and thiol chemistry (1-3). RSS are characterized by sulfane sulfur– containing molecules, including per- and polysulfides, in which sulfur exists in the zero-oxidation state (S^0^) (4, 5). Sulfane sulfur species can spontaneously undergo sulfur redistribution and condensation to form elemental sulfur, predominantly as the cyclo-octasulfur (S_8_) allotrope, representing the thermodynamic endpoint of sulfane sulfur chemistry (6, 7). Given the high abundance of RSS in many biological systems, including bacteria and mammals, S_8_ is predicted to be generated endogenously as a terminal product of sulfane sulfur metabolism (1, 4, 8, 9). Detection and quantification of biological S_8_ remains challenging due to its low relative abundance, insolubility in aqueous media, and dynamic equilibrium with other sulfane sulfur species (10, 11). Elemental sulfur can be structurally identified by Raman spectroscopy, however, its utility in biological samples remains limited due to low sensitivity and specialized instrumentation (12, 13). Pt-macrocyclic host–guest capture of S_8_ followed by LC–MS detection, provides an elegant molecular recognition of elemental sulfur (14); however, this approach relies on bespoke metal– organic host architectures that are not widely accessible, restricting its general applicability and routine use. As a result, information on elemental sulfur (S^0^) in biological samples remains limited, despite chemical arguments for its formation. Here, we describe a chemical method for selectively measuring elemental sulfur and selenium in biological matrices using broadly accessible reagents and techniques. Using complementary spectroscopic and mass-spectrometric readouts, we demonstrate selective isolation, orthogonal validation, and quantitative analysis of zero-valent chalcogen species across diverse sample types.

## Material and Methods

### Reagents

Elemental sulfur (S_8_), cystine, sodium selenite (Na_2_SeO_3_), triphenylphosphine (TPP), and carbon disulfide (CS_2_) were purchased from Sigma-Aldrich and used without further purification. Deutero-chloroform (CDCl_3_) for NMR analysis was obtained from Cambridge Isotope Laboratories. Phosphate-buffered saline (PBS) was purchased from Gibco. Protein concentrations were determined using the bicinchoninic acid (BCA) assay kit (Pierce) according to the manufacturer’s instructions.

### Bacterial culture

*E coli* (BW25113) wild-type strain transformed with an ampicillin resistance-containing plasmid were cultured on LB agar plates with ampicillin (100ug/mL). Single colonies cultured in LB media with ampicillin (100ug/mL) at 37C, 250 rpm to stationary phase. Cultures were diluted to OD=0.4-0.6 (∼1:100 dilution) and cultured in LB media supplemented with cystine or sodium selenite for 48 hours as described (15).

### Human cancer cell line

HeLa cells were cultured in Dulbecco’s Modified Eagle Medium (DMEM; high glucose) supplemented with 10% (v/v) fetal bovine serum (FBS). Cells were maintained at 37 °C in a humidified atmosphere containing 5% CO_2_. Culture medium was replaced every 2–3 days and cells were passaged at ∼80% confluence using trypsin-EDTA. Media was supplemented with 1 mM cysteine for 24 hours prior to S_8_ quantification.

### Human tumour samples

Human tumour explants were obtained from patients undergoing surgical resection at Leicester Cancer Research Centre with approval from Health Research Authority (study number 174618, ethics number 15/YH/0510). All participants provided written informed consent. Samples were anonymised prior to analysis and all procedures were conducted in accordance with the Declaration of Helsinki. Tumour explants were cultured at the liquid-air interface in DMEM-F12 media supplemented with autologous serum for 24 hours as previously described (16, 17), prior to S_8_ measurements.

### Elemental Sulfur/Selenium extraction

Samples were sonicated in cold PBS on ice. Liquid samples (eg, conditioned media) were diluted in equal volume buffer. Biological samples were extracted with equal volumes of CS_2_ by vortexing for 12 cycles of 30 seconds. Samples were then centrifuged (5000 x*g*, 3 minutes) to separate aqueous and organic layers.

### Mouse Tissue Collection and Sample Preparation

Mouse faeces were collected under University of Leicester-approved ethical protocols from C57BL/6 mice (>12 weeks old) under AWERB approval [AWERB_2025_239]. Faecal samples were hydrated overnight at 4C and homogenised in phosphate-buffered saline (PBS) via sonication. Homogenised samples were subjected to biphasic extraction with carbon disulfide (CS_2_) as described above, followed by derivatization with TPP for GC-MS quantification of elemental sulfur.

### TPP derivatisation and NMR quantification

Extracted S8 is reacted with 1/10^th^ volume of 100 mM TPP dissolved in CS_2_ at room temperature for 1 hour. For NMR quantification, CDCl3 is added to the solution (25% v/v final) and NMR spectra acquired. NMR spectra were acquired on a spectrometer operating at 242.94 MHz for ^31^P nuclei at 298 K. A 90° pulse (P1 = 13 µs) was applied at a power level of 45 W, with a receiver gain of 2050. The relaxation delay (D1) was 2.0 s, and acquisition parameters included a dwell time of 6.8 µs and a decoupling delay of 7.68 µs.

Spectra were processed using standard exponential multiplication and baseline correction. For quantitative analysis, the integration of the phosphine sulfide peak was used to determine elemental sulfur content from an authentic S_8_-generated standard curve. NMR spectra were processed and analysed using Spectrus software (ACD/Labs), including baseline correction and peak integration. Peak integrals were used for quantification of derivatized products, with integration thresholds and baseline correction parameters kept consistent across samples.

### GC-MS quantification

GC-MS analysis was carried out using an Agilent 8890 GC (Agilent Technologies, Wokingham, UK) and LECO Pegasus BT 4D time-of-flight MS (LECO Instruments UK Ltd., Cheshire, UK) with PAL3 autosampler (CTC Analytics, Zwingen, Switzerland). A DB-5MS capillary column (l = 20 m, I.D. = 0.18 mm, dF = 0.18 μm from Agilent Technologies) was used for the analysis. The GC conditions were as follows: the column starting temperature was 40 °C, which was raised to 325 °C at 15 °C/min where it was held for 3 minutes giving a total run time of 22 min. The inlet temperature was maintained at 280 °C with a septum purge flow of 3 mL/min. The GC was operated in splitless mode with a purge activation time of 30 s and purge flow of 20 mL/min, with 1.5 mL/min column flow rate using helium as a carrier gas (N6.0, BOC, Woking, UK). Samples were prepared in chloroform and 1 µl was injected. The mass spectrometer was operated in electron ionisation (EI) mode at 70 eV, with a scan range of 50 to 400 amu, an extraction frequency of 26 kHz, an acquisition rate of 10 spectra/s, and a solvent delay of 300 s. The transfer line to the mass spectrometer was heated to 280 °C and the source temperature was maintained at 300 °C. The TPP=S adduct eluted at a retention time of 15.85 min.

Data analysis was performed using LECO ChromaTOF software. Target analyte finding was used to identify the TPP=S peak based on the extracted ion chromatogram at m/z 183, with m/z 262 and m/z 294 used as qualifier ions. Peak identity was confirmed by matching to the retention time of authentic standards prepared at concentrations of 5–50,000 nM. Peaks were accepted where a signal-to-noise ratio of >10 was achieved. Peak areas were exported from ChromaTOF and a multipoint calibration curve was constructed, from which sample concentrations were calculated. Three replicates were analysed per condition.

### Raman spectroscopy

Carbon disulfide (CS_2_) solutions containing either authentic cyclo-octasulfur (S_8_) or CS_2_ extracts derived from cellular samples were deposited onto Raman-compatible glass microscope slides (MirrIR slides; Kevley Technologies, Parma, Ohio) and allowed to air dry completely at ambient temperature prior to analysis. Raman spectra were acquired using an AIRsight Raman/IR microscope (Shimadzu) equipped with a 532 nm excitation laser, operated under confocal Raman conditions using low laser power to minimize photothermal effects. For each sample, spectra were collected from 5–10 distinct positions across the dried deposit to account for spatial heterogeneity and averaged prior to analysis.

### Selenite reactions

Sodium selenite (1 mM, final concentration) was dissolved into PBS and either sodium borohydride (approx. 50 mM, final concentration) or sodium chloride (50 mM) was added and vortexed. The solutions were then extracted with CS_2_ containing 10 mM TPP. CDCl_3_ was added (25% v/v) and samples analysed by ^31^P-NMR.

### Statistical analyses

Data are presented as mean ± standard error of the mean (SEM). Statistical analyses were performed using GraphPad Prism 10, including linear regression, unpaired two-tailed t tests, and one-way ANOVA with post hoc multiple comparison tests where appropriate. A p-value < 0.05 was considered statistically significant.

## Results

### Chemical Basis and Workflow for Selective Elemental Sulfur Quantification

To develop a chemical assay for the selective isolation and quantification of S^0^, its physicochemical properties and its reactivity toward triorganyl phosphines were utilized (7, 15, 18). The S_8_ allotrope of S^0^ is solubilized by carbon disulfide (CS_2_), enabling physical separation from aqueous sulfur species and biological nucleophiles (Figure 1). The extracted S_8_ is subsequently derivatized with triphenylphosphine (TPP) to form triphenylphosphine sulfide (TPP=S) (15), which is quantified by ^31^P−NMR or GC-MS. The extraction process re-moves interfering aqueous-soluble sulfur species (including protein and small molecule thiols, per- and polysulfides, thiosulfate, etc.) from S_8_ prior to derivatization. Accordingly, the method is designed to quantify neutral, zero-valent chalcogen allotropes that are extractable into non-polar organic solvent, with S_8_ as the prototypical sulfur species examined in this study.

**Figure 1.**
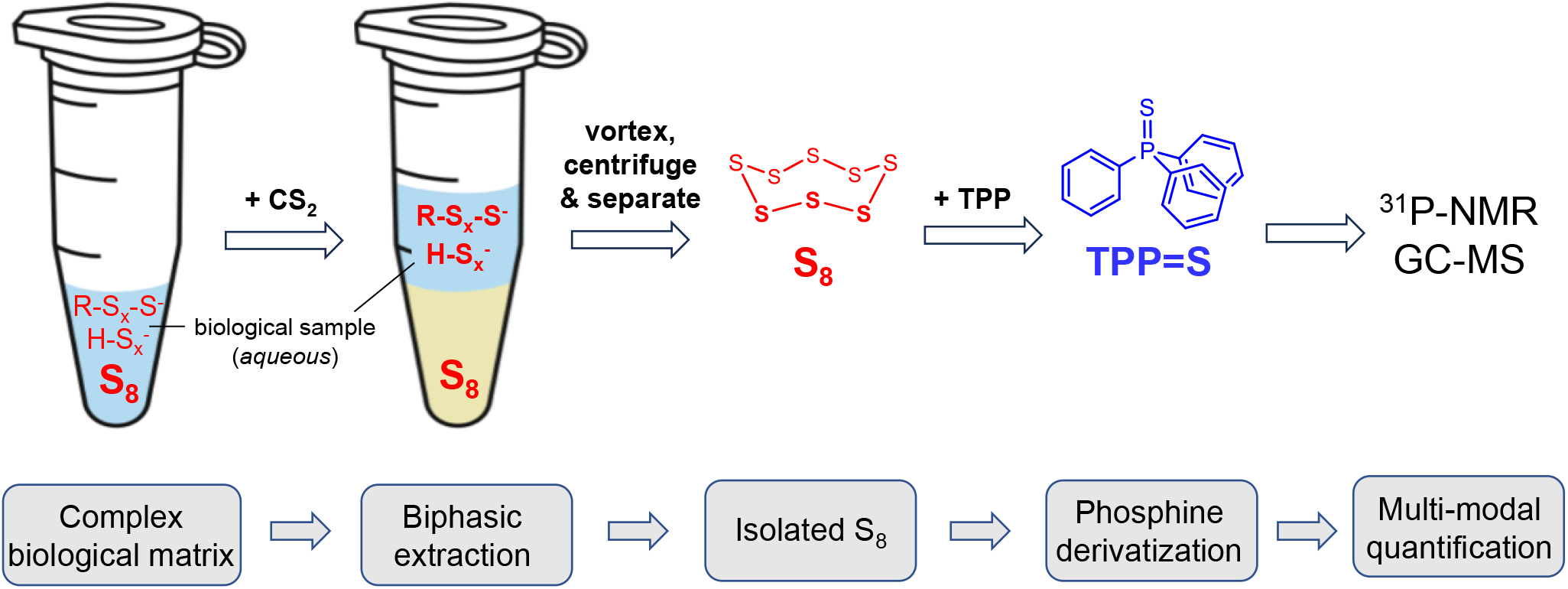
Workflow for extraction and quantification of elemental sulfur (S_8_) from biological samples. Aqueous biological samples are homogenised and extracted with carbon disulfide (CS_2_), which selectively solubilises elemental sulfur. Following centrifugation, the CS_2_ layer is isolated and reacted with triphenylphosphine (TPP) to form triphenylphosphine sulfide (TPP=S). The derivatised product is then quantified using either ^31^P-NMR spectroscopy or GC–MS.

### Stoichiometric Derivatization of Elemental Sulfur by Triphenylphosphine

To establish the linearity of S_8_ detection by phosphine derivatization, known amounts of elemental sulfur were dissolved into CS_2_ and reacted with excess TPP. The resulting TPP=S formation was quantified to assess stoichiometric conversion. Using ^31^P−NMR spectroscopy (Figure 2A), formation of TPP=S exhibited a linear dependence on input S_8_ concentration across the micromolar range examined (R^2^= 0.994) (Figure 2B), consistent with stoichiometric derivatization of S^0^. To extend the analytical range, GC–MS detection of the TPP=S adduct was used. GC–MS analysis of the TPP=S adduct exhibited a linear response to S_8_ concentration across the nanomolar range examined when plotted on logarithmic axes (R^2^= 0.984) (Figure 2C), with linearity maintained down to 5 nM, the lowest concentration within the calibrated range. Therefore, linearity was established within the respective operating ranges of ^31^P−NMR (micromolar) and GC–MS (nanomolar), together defining a broad quantitative window for elemental sulfur analysis.

**Figure 2.**
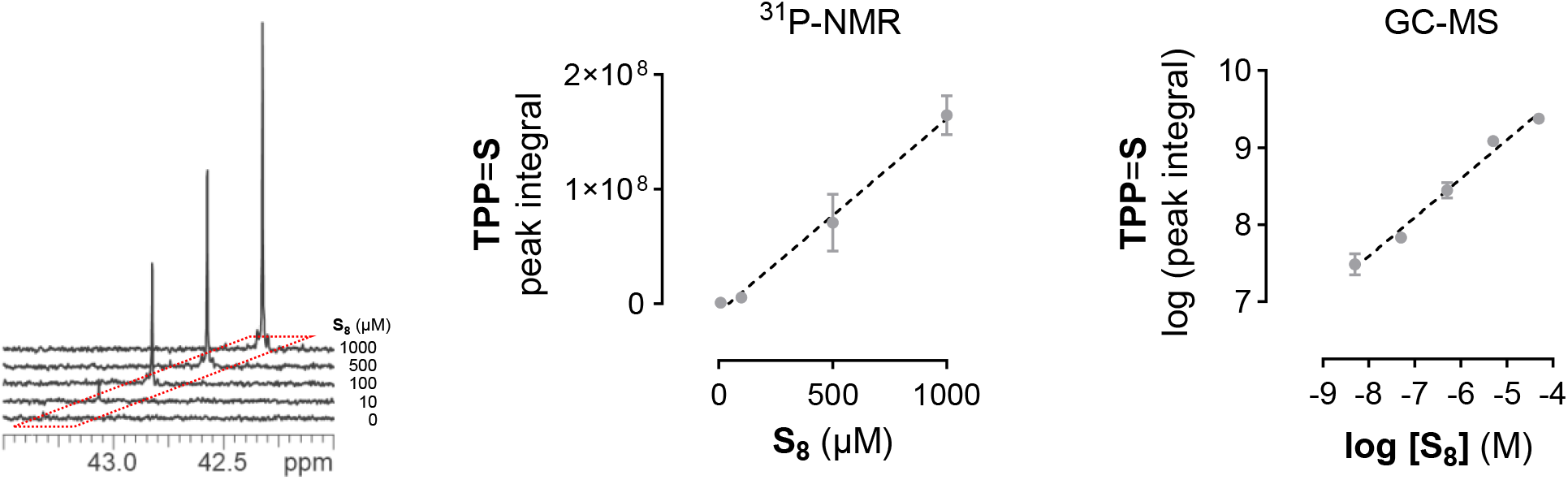
Derivatization of elemental sulfur (S_8_) with triphenylphosphine (TPP) and quantification via ^31^P-NMR or GC-MS. (A) Representative ^31^P-NMR spectra of triphenylphosphine sulfide (TPP=S, +43.3 ppm) formed by reacting authentic S_8_ with 10 mM TPP in CS_2_/CDCl_3_. (B) Calibration curve showing linear relationship between S_8_ concentration and TPP=S NMR peak integral values (au). (C) Log-log plot of TPP=S integrated peak area compared to S8 concentration showing linear relationship. Data in (B) and (C) are presented as mean ± SEM (n = 3) and linear regression analysis was performed using GraphPad Prism 10; dashed line indicates best-fit regression.

### Effect of Nucleophiles on Phase-Selective Isolation of Elemental Sulfur

S_8_ is electrophilic and reactive with thiols (6, 19, 20). To determine whether aqueous thiols interfere with elemental sulfur isolation and subsequent detection, S_8_ was extracted in the presence of cysteine prior to phosphine derivatization. The presence of cysteine during ex-traction did not measurably alter the recovered TPP=S signal relative to S_8_ alone (Figure 3A). These data indicate that cysteine does not intercept or consume elemental sulfur under the extraction conditions employed. The absence of signal attenuation supports a phase-selective mechanism in which S_8_ is physically partitioned into the CS_2_ phase prior to derivatization, rendering it inaccessible to aqueous-phase thiols. Thus, competing aqueous nucleophiles do not interfere with S_8_ isolation or quantification within the defined work-flow.

**Figure 3.**
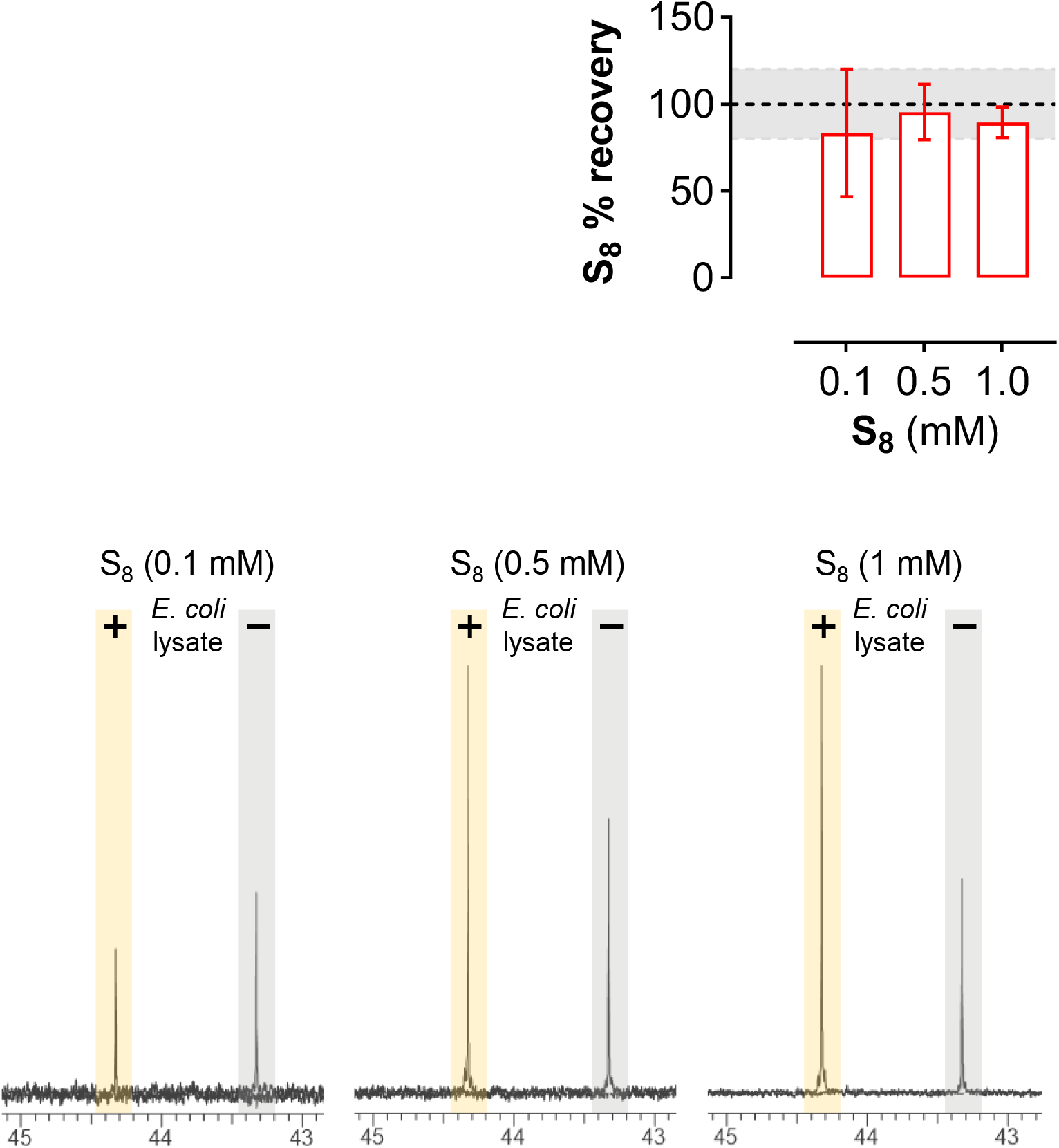
Influence of biological matrix on S_8_ recovery and TPP=S quantification. (A) Bar graph showing TPP=S peak integrals following extraction of authentic S_8_ in the presence or absence of cysteine. Cysteine was added to the aqueous phase prior to CS_2_ extraction. Data are presented as mean ± SEM (n = 3). (B) Bar graph showing calculated S_8_ recovery (%) from *E. coli* lysates compared to PBS control. Data are mean ± SEM (n = 3). The shaded region (80–120%) indicates efficient analyte recovery. (C) Representative ^31^P-NMR spectra of TPP=S (+43.3 ppm) derived from S_8_ spiked into either PBS or *E. coli* lysate. Spectra are overlaid and offset by +1.0 ppm for clarity.

### Matrix Effects on Elemental Sulfur Recovery in Whole Cell Lysate

Having established that aqueous thiols do not interfere with phase selective isolation of elemental sulfur, we next evaluated the effect of a complex biological matrix on S_8_ recovery and detection. Whole cell *E. coli* lysate was chosen as a representative protein and metabolite rich environment containing a broad distribution of potential nucleophiles. Elemental sulfur was introduced into clarified *E. coli* lysate prior to CS_2_ extraction and phosphine derivatization, and recovered TPP=S was quantified under identical conditions to buffer controls (Figure 3B). S_8_ remained readily detectable following extraction from lysate, although absolute recovery was reduced relative to buffer alone (Figure 3B). This attenuation was concentration dependent and reproducible across independent preparations, indicating a matrix associated effect on recovery rather than loss of analytical selectivity. No additional phosphorus containing species were observed by ^31^P−NMR or GC–MS analysis, confirming that lysate components do not generate interfering signals or chemically distinct derivatization products. Together, these data demonstrate that while interactions with complex cellular matrices can influence extraction efficiency, phase selective isolation preserves chemical specificity for elemental sulfur and enables quantitative detection within a defined analytical range.

### Orthogonal Spectroscopic Validation of Extracted Elemental Sulfur

Raman spectroscopy was employed to orthogonally verify the chemical identity of the sulfur species detected by TPP derivatization, as this technique enables direct identification of S_8_ without chemical modification. Escherichia coli cultures supplemented with cystine were used as a reproducible cellular system in which RSS accumulation occurs in a cystine-dependent manner (15). CS_2_ extracts exhibited Raman features characteristic of S_8_, in agreement with authentic standards (Figure 4A), confirming elemental sulfur as the extracted analyte.

**Figure 4.**
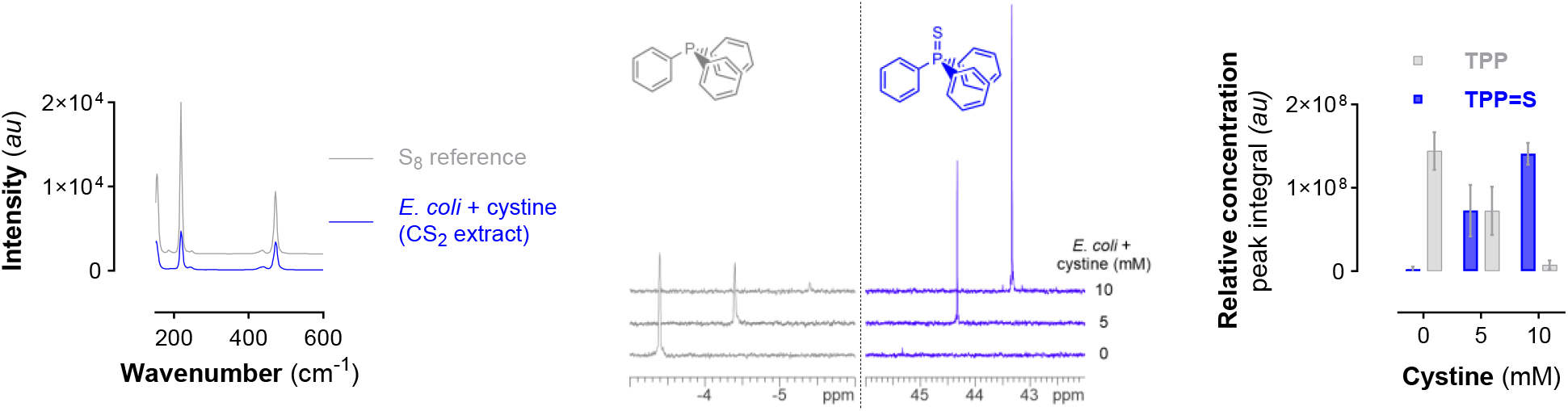
Detection of S_8_ in hypoxic *E. coli* cultures supplemented with cystine. (A) Raman spectra of authentic S_8_ reference or CS_2_ extract of hypoxic *E. coli* cultured with 5 mM cystine for 44 hours (B) Representative ^31^P-NMR spectra showing triphenylphosphine (TPP) and triphenylphosphine sulfide (TPP=S) signals following derivatisation of CS_2_ extracts from hypoxic *E. coli* cultures. Spectra are offset by +1.0 ppm for clarity. (C) Linear increase of TPP=S peak integrals (*au*) and decrease of unreacted TPP from *E. coli* cultures grown with cystine. Data are presented as mean ± SEM (n = 3).

Quantitative assessment of S_8_ accumulation in *E. coli* as a function of cystine concentration was subsequently measured. A cystine-dependent increase in extracted S_8_ was observed (Figure 4B-C). Quantification using the ^31^P−NMR standard curve (Figure 2B) yielded S_8_ concentrations of 475 µM and 880 µM (corresponding to 285 nmol and 528 nmol) in response to cystine supplementation corresponding to 5 mM and 10 mM nominal cystine equivalents, respectively. Together, Raman spectroscopy confirms the presence of S_8_, while ^31^P−NMR quantification demonstrates a cystine-dependent increase in extracted elemental sulfur from *E. coli* cultures.

### Applicability of the Extraction and Quantification Protocol across Biological Matrices

To evaluate the robustness and general applicability of the extraction and quantification workflow, we applied the protocol to chemically distinct biological matrix types, encompassing liquid, solid, cellular, and tissue samples. In *E. coli* conditioned media, S_8_ was readily detected only following supplementation with high concentrations of cystine (nominally 10 mM) under hypoxic/anaerobic conditions, reaching ∼100–120 nmol/mL, whereas negligible signal was observed in the absence of cystine supplementation (Figure 5A). Patient-derived head and neck (HN) or mesothelioma (TH) tumour explants cultured under normoxic conditions exhibited relatively low S_8_ levels (∼30–70 pmol/mg protein) (Figure 5B), consistent with variable metabolic and diffusional constraints in complex, heterogeneous tissues, whereas an *in vitro* cellular monoculture (HeLa) showed higher levels (∼1 nmol/mg protein) (Figure 5C). In solid samples (mouse faecal pellets), S_8_ was detected at ∼3–6 nmol per pellet (Figure 5D), reflecting the heterogeneous and microbially enriched nature of this matrix. Although there is currently no established baseline for S_8_ concentrations in biological systems, the measured values are internally consistent across matrices and follow a logical hierarchy, with the highest levels observed in extracellular, sulfur-rich conditions and lower levels in structured cellular and tissue environments. These data therefore provide a first quantitative framework for S_8_ in biology and demonstrate that the method yields chemically and biologically plausible measurements across liquid, cellular, tissue, and solid samples.

**Figure 5.**
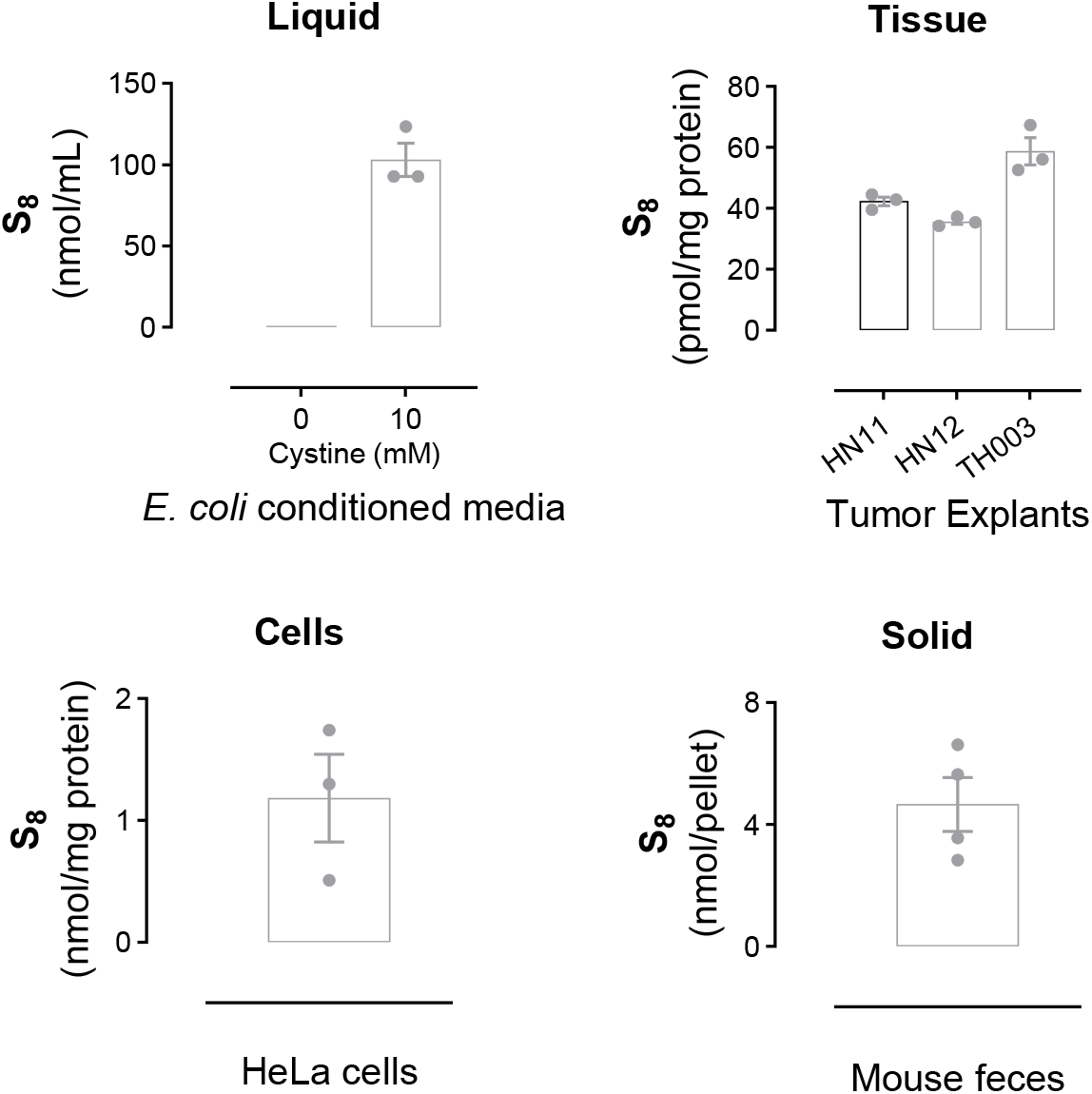
Detection of elemental sulfur (S_8_) across diverse biological matrices. Elemental sulfur (S_8_) was quantified in multiple biological systems using CS_2_ extraction followed by triphenylphosphine (TPP) derivatization and ^31^P-NMR analysis. (A) S_8_ levels detected via ^31^P-NMR in conditioned media from hypoxic *E. coli* cultured with or without 10 mM cystine, expressed as ng/mL media. (B) S_8_ detected in human tumour explant samples, normalised to protein content. (C) S_8_ measured in human carcinoma cell line MDA-MB-134, normalised to protein content, with individual data points representing independent experiments. (D) S_8_ detected in mouse faecal pellets, expressed as nmol/pellet. Data are shown as individual biological replicates.

### Extension of the Workflow to Elemental Selenium

To extend the workflow to selenium redox chemistry, we investigated the formation and detection of elemental selenium (Se^0^) using TPP derivatization and ^31^P−NMR spectroscopy. Initial experiments confirmed that selenite (SeO_3_^2−^) in buffer does not react with TPP, consistent with its fully oxidized oxidation state. In contrast, chemical re-duction of selenite using sodium borohydride (BH_4_^−^) generated a distinct ^31^P−NMR resonance corresponding to triphenylphosphine selenide (TPP=Se) at +35.3 ppm (15), demonstrating selective trapping of reduced selenium by TPP (Figure 6A–B).

**Figure 6.**
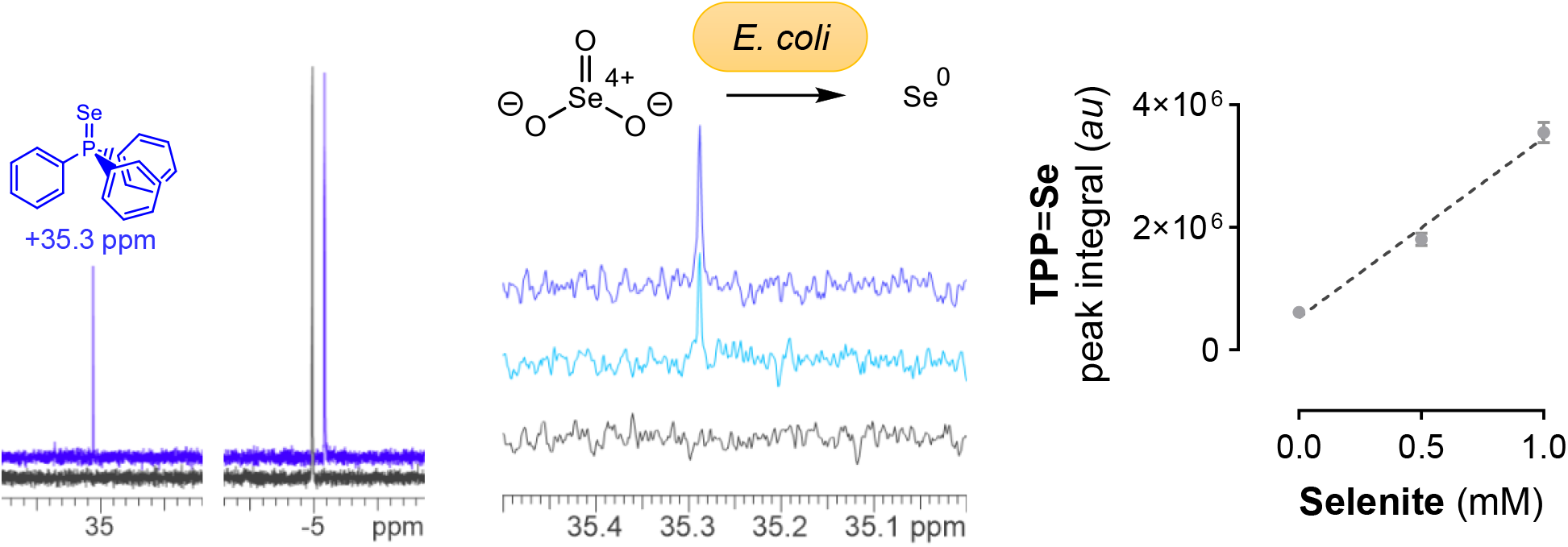
Detection and quantification of elemental selenium (Se^0^) via TPP derivatisation and ^31^P-NMR. (A) Representative ^31^P-NMR spectra of CS_2_ extracts from PBS containing either sodium selenite alone (black) or selenite reduced with sodium borohydride (blue). The peak at +35.5 ppm corresponds to triphenylphosphine selenide (TPP=Se) (15). Spectra are offset by –0.5 ppm for clarity. (B) Schematic illustrating the reduction of selenite to elemental selenium (Se^0^) by sodium borohydride, followed by derivatisation with TPP to form TPP=Se. (C) Representative ^31^P-NMR spectra of TPP=Se formed from CS_2_ extracts of *E. coli* cultured with selenite. Control (black), 0.5 mM selenite (light blue), and 1.0 mM selenite (dark blue) conditions are shown. (D) Quantification of TPP=Se peak integrals as a function of selenite concentration. Data are presented as mean ± SEM (n = 3). Linear regression analysis was performed using GraphPad Prism 10; dashed line indicates best-fit regression.

Building on this chemical validation, *E. coli* cultures were supplemented with selenite (0, 0.5, and 1 mM), followed by CS_2_ extraction and TPP derivatization. ^31^P−NMR analysis of the resulting extracts revealed a selenite-dependent in-crease in the TPP=Se signal (Figure 6C), indicating cellular reduction of selenite to elemental selenium under these conditions. Together, these results demonstrate that the extraction and derivatization workflow is applicable to zero-valent selenium and can be used to detect and quantify Se^0^ formation in biological samples.

## Discussion

Elemental sulfur represents a thermodynamically stable endpoint of RSS chemistry but has remained difficult to measure directly in biological systems. The work presented here establishes a chemically selective and broadly accessible workflow that enables direct detection and quantification of zero-valent sulfur, addressing a longstanding analytical gap in sulfur biology (21-23). Selectivity in this approach arises from the intrinsic physicochemical properties of cyclo-octasulfur. Phase-selective extraction physically separates elemental sulfur from aqueous sulfur species prior to derivatization, while stoichiometric reaction with triphenylphosphine affords a chemically defined and spectroscopically tractable readout. This strategy decouples S_8_ measurement from the broader RSS pool, allowing elemental sulfur to be distinguished from thiols, persulfides, polysulfides, and other sulfane sulfur species that confound indirect or nucleophile-based assays.

Quantitative performance across a wide dynamic range further enhances the utility of the method. Stoichiometric derivatization monitored by ^31^P−NMR provides a calibration-independent readout in the micromolar to millimolar range, while GC–MS analysis of the same phosphine adduct extends sensitivity into the nanomolar range. Together, these complementary modalities define a practical quantitative window spanning multiple orders of magnitude without altering the underlying chemistry.

The workflow was compatible with chemically diverse biological matrices, including conditioned media, cells, tissues, and solid biological material, without matrix-specific re-optimization. Variability in recovered elemental sulfur reflected expected matrix complexity rather than loss of chemical selectivity, underscoring the robustness of the extraction-derivatization strategy. Importantly, the method relies exclusively on widely available reagents and standard analytical instrumentation, facilitating routine implementation and broad adoption across laboratories. Extension of the workflow to elemental selenium demonstrates its generality toward zero-valent chalcogen species. Selective trapping of reduced selenium, but not oxidized selenite, confirms that the approach is oxidation-state zero chalcogen selective rather than sulfur-specific.

To place these measurements in a cellular context, S_8_ levels determined in HeLa cells (∼1–1.5 nmol/mg protein) can be converted to apparent intracellular concentrations by assuming a typical cellular protein content of ∼200–300 mg/mL. This yields an estimated S_8_ equivalent of ∼200–450 µM. While this value is higher than might be intuitively expected for a low-abundance redox-active species, it should be emphasised that this represents total extractable S_8_ rather than freely diffusible sulfur. Given the hydrophobic nature of S_8_, substantial partitioning into membranes, lipid environments, or protein-associated pools is likely, such that the measured values reflect compartmentalised or sequestered sulfur rather than a uniform cytosolic concentration. In this context, the observed levels are chemically plausible and consistent with the capacity of cells to accumulate and stabilise nonpolar sulfur species. These findings therefore suggest that S_8_ may constitute a significant and previously underappreciated intracellular sulfur reservoir.

Although the present workflow relies on chemical derivatization, it is distinct from many indirect approaches used previously to interrogate sulfur species in biology (21, 23). In this case, derivatization occurs only after physical isolation of elemental sulfur and proceeds via a stoichiometric, oxidation-state-specific reaction, yielding a single, chemically defined product. Consequently, the derivatization step reports directly on zero-valent sulfur itself rather than on downstream reactivity, secondary metabolites, or proxy species. The combination of phase-selective isolation and chemically specific derivatization therefore enables indirect measurement in an analytical sense, while retaining chemical unambiguity and selectivity for elemental sulfur. Importantly, this strategy is implemented using widely available reagents and standard analytical platforms, offering a level of accessibility and transferability that is difficult to achieve with receptor-based or probe-dependent methods.

Overall, this work transforms elemental sulfur and selenium from chemically inferred endpoints of RSS metabolism into directly measurable species. By providing a selective, accessible, and quantitative analytical strategy, the approach establishes a foundation for routine interrogation of zero-valent chalcogen chemistry in biological systems.

## Conclusion

We present a chemically selective and broadly accessible strategy for the detection and quantification of elemental sulfur and selenium in biological matrices. By combining phase-selective isolation with stoichiometric phosphine derivatization and complementary spectroscopic readouts, this approach enables direct measurement of zero-valent chalcogen species across diverse sample types. The work-flow provides a practical foundation for routine interrogation of elemental sulfur and selenium in biological systems and removes a key analytical barrier in Reactive Sulfur/Selenium Species research

## Acknowledgements

The authors thank Dr Gareth Miles, and Prof Catrin Pritchard (Leicester Cancer Research Cenre) for providing patient-derived tumor explant samples and Dr. Ralph Vokes (Shimadzu UK) for assistance with Raman spectroscopy measurements. This research was supported by The Royal Society (RG\R1\241248).

## CRediT authorship contribution statement

**Christopher H. Switzer:** Conceptualization, Methodology, Investigation, Formal analysis, Funding acquisition, Writing – original draft, Supervision. **Nada E. Eldakra:** Investigation, Formal analysis, Validation, Writing – review & editing. **Ethan J. York:** Investigation, Validation. **Rebecca L. Cordell:** Methodology, Resources (GC–MS). All authors reviewed and approved the final manuscript.

## Data Statement

All raw and processed data supporting the findings of this study, including NMR spectra, GC–MS data, and associated analytical files, have been deposited in the University of Leicester Figshare repository (figshare.le.ac.uk) under DOI: 10.6084/m9.figshare.xxxxxxx.

